# Established Sulfopeptide Tandem Mass Spectrometry Behavior and Sulfotransferase Assays Refute Tyrosine Sulfation as a Histone Mark

**DOI:** 10.1101/2024.09.23.613127

**Authors:** Menatallah M. Youssef, Carson W. Szot, Miriam F. Ayad, Lobna A. Hussein, Maha F. Abdel-Ghany, Ryan C. Bailey, Ahmed M. Mostafa, Kristina Hakansson

## Abstract

Tyrosylprotein sulfotransferases (TPST1 and TPST2) are only known to exist in the Golgi^1^. Thus, Yu et al.’s findings that histone H3 sulfation occurs in the cytosol are surprising. Cytosolic sulfotransferase SULT1B1, which sulfates phenolic small molecules, was proposed to catalyze H3Y99 sulfation with 3′-phosphoadenosine 5′-phosphosulfate (PAPS) as SO_3_ donor. Bottom-up liquid chromatography-tandem mass spectrometry (LC-MS/MS)-based proteomic analyses were key to this conclusion. Such experiments appear to have been mainly performed following standard protocols, including reduction (reagent unknown), alkylation with iodoacetamide (IAA), and proteolytic digestion prior to nanoflow LC (nanoLC) separation of proteolytic peptides and data-dependent higher energy collision dissociation (HCD) to generate sequence-informative peptide fragments. However, the published HCD MS/MS spectra do not match established sulfopeptide fragmentation behavior. We present reannotation of these spectra and additional experiments that found a lack of evidence to support H3Y99 sulfation.

---

One not immediately obvious protocol modification in Yu et al. appears to involve double digestion with trypsin and Glu-C to generate the proteolytic peptide AyLVGLFEDTNLC(carbamidomethyl)AIHAK (“y” indicates the proposed Y99sulf). While this enzyme combination is not directly mentioned, nor motivated, evidence is provided in Supplementary Table 1 (Yu et al.), which indicates two missed cleavages (presumably following E105 and D106). Additionally, recombinant human (rh) histone H3.2 (New England BioLabs; Uniprot entry Q71DI3) includes E97, translating into the peptide starting at A98. Double digestion is warranted as trypsin alone would generate the longer peptide FQSSAVMALQEASEAyLVGLFEDTNLC(carbamidomethyl)AIHAK), which may be more difficult to identify. However, it is unclear why double digestion was chosen *before* the Y99 modification site was found.

Interestingly, MS1 signals (2+ charge state, ∼m/z 1058; Yu et al, Figure 1c) corresponding to the shorter proteolytic peptide with monoisotopic mass shifts +79.9583 and +79.9666 were detected from nuclear proteins at two different LC retention times (86.3 vs. 81.7 minutes; Yu et al, Figure 1b). These mass shifts suggest both H3 sulfation and phosphorylation in the nucleus with phosphorylation being a known histone modification^2^. HCD MS/MS spectra for the presumed sulfo- and phosphopeptide are shown in Yu et al., Figures 1a and Extended Data Figure 1a. **The annotated sulfopeptide MS/MS spectrum (Yu et al**., **Fig. 1a, and Figure 1a here) caught our eye as it does not reflect known sulfopeptide fragmentation behavior**. First, it is well established that tyrosine sulfation is extremely labile in the gas phase^3-8^. Typically, sulfate-retaining fragment ions are entirely absent following collisional activation such as HCD. Yet, the presumed sulfopeptide HCD MS/MS spectra (Yu et al., Fig. 1a and Extended Data Fig. 2g) show 100% sulfate retention, which would be remarkable. For example, all annotated N-terminal *b*-type ions in Fig. 1a contain the proposed sulfotyrosine with no hint of even partial desulfation. In addition, it appears from the sequence map (Yu et al., Figure 1a) that the larger, C-terminal *y*_16_^2+^ fragment contains the sulfotyrosine. However, this label is in the wrong location: the *y*_16_ fragment contains the 16 C-terminal residues, i.e., backbone cleavage occurred between the tyrosine and leucine, not between the alanine and tyrosine, which would generate a *y*_17_ fragment (not annotated). Moreover, the reported m/z values for the annotated sulfopeptide fragments (Yu et al., Fig. 1a) more accurately match the expected theoretical fragments for the tyrosine <u>phospho</u>peptide: the mass errors for <u>phosphate</u>-retaining *b*-type fragments are less than 3 ppm, whereas they range from 12.5 to 31.5 ppm for the annotated sulfate-retaining *b* ions (**Extended Data, Table 1**).

**Figure 1.**
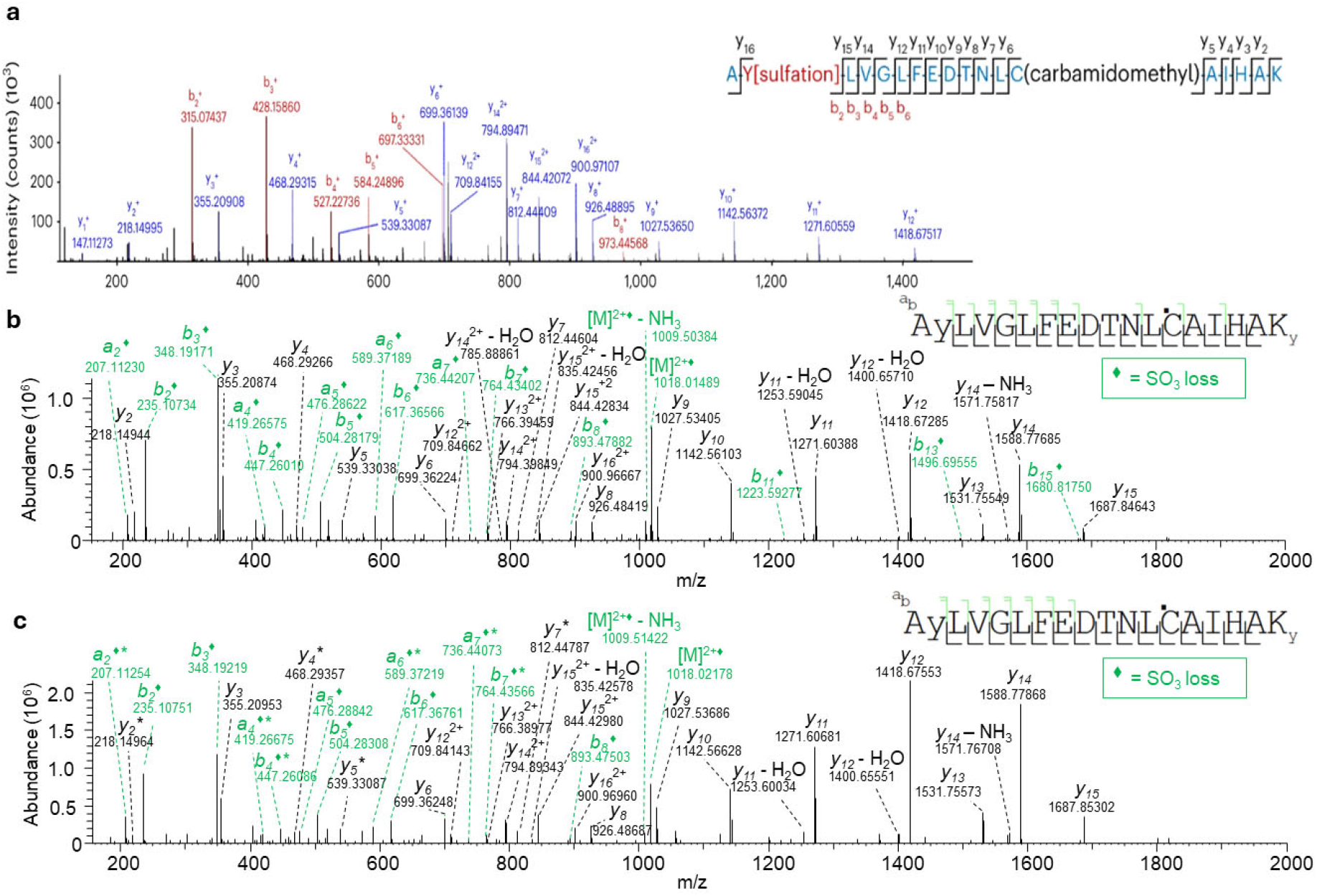
H3Y99 sulfation is misannotated in Yu et al. HCD MS/MS spectra. **a**, HCD MS/MS spectrum of proposed AyLVGLFEDTNLC(carbamidomethyl)AIHAK, (y indicates sulfotyrosine), acquired and annotated by Yu et al., showing *b*-type ions retaining the highly labile sulfate modification and misplaced *y*_14_-*y*_16_ labels. Annotated *b*-type fragments more accurately align with a phosphopeptide assignment. **b**, HCD MS/MS spectrum of synthetic AyLVGLFEDTNLC AIHAK (indicates carbamidomethylated cysteine), exhibiting expected complete SO_3_ neutral loss (highlighted in green) from sulfate-containing fragments. **c**, HCD MS/MS spectrum observed at 87.58 min upon reexamination of Yu et al.’s raw data from a 120 min gradient. In contrast to the spectrum in (a), reassigned as a phosphopeptide, the spectrum in (c) displays known sulfopeptide fragmentation behavior with complete loss of the sulfate PTM and is highly similar to the synthetic sulfopeptide spectrum in (b).

To further refute the sulfopeptide annotation (Yu et al., Fig. 1a), we obtained the synthetic sulfopeptide AyLVGLFEDTNLCAIHAK from Genscript (Piscataway, NJ) and subjected it to reduction (with dithiothreitol), alkylation (with IAA), and nanoLC-HCD MS/MS on the same instrument type used by Yu et al. (Orbitrap Fusion Lumos). As anticipated, the corresponding HCD spectrum (**our Figure 1b**) showed a complete absence of sulfate-retaining fragments. We also reexamined the raw nanoLC-HCD MS/MS data (PRIDE accession ID PXD043754) from which Yu et al.’s Fig. 1a was generated. This reanalysis identified an HCD spectrum (**Figure 1c**) with a fragmentation pattern similar to our synthetic sulfopeptide counterpart, suggesting that histone sulfation may still be present.

Yu et al. also acquired the synthetic sulfopeptide and subjected it to nanoLC-HCD MS/MS analysis under different LC conditions (shorter gradient) than their aforementioned proteomic analysis. However, the presented MS/MS spectrum (Yu et al., Fig. 1e, bottom) at an elution time ∼14.5 min (Yu et al., Fig. 1d, bottom) again does not match known sulfopeptide HCD fragmentation behavior. Short gradient nanoLC-HCD MS/MS analysis was also performed for a HepG2 nuclear extract with and without addition of the synthetic sulfopeptide, again showing HCD spectra (Yu et al, Figs. 1e, top, middle) mismatched with a sulfopeptide annotation. Upon reexamination of the corresponding raw data, the H3Y99 <u>phospho</u>peptide was observed to elute at 14.09-14.18 minutes in the three datasets (**Extended Data, Figure 1, left**) as confirmed by the corresponding HCD spectra. By contrast, Yu et al. reported phosphopeptide elution ∼25.5 min (Yu et al., Extended Data, Fig. 1b). However, while a peptide with similar m/z ratio is present in their nuclear extract data at 24.45 min, the corresponding abundance (∼2.56×10^3^) is significantly lower than the phosphopeptide abundance at 14.09 min. (∼1.56×10^7^; **Extended Data, Figure 1**). Also, there is no HCD spectrum for the latter precursor ion. For the sulfopeptide and mixed samples, there is no signal ∼25.5 min. On the other hand, HCD spectra corresponding to a H3Y99 <u>sulfo</u>peptide are present for all three samples ∼14.6 min (**Extended Data, Fig. 2**), i.e., the phospho- and sulfopeptide are eluting in the same order as for the longer LC gradient.

Several observations from our reexamination of these raw data are puzzling, e.g., why the histone <u>phospho</u>peptide is observed when the synthetic <u>sulfo</u>peptide was injected (Extended Data, Fig. 1). We subjected the shorter gradient LC-HCD MS/MS data to bioinformatic analysis (Proteome Discoverer) and noted significant carryover (Extended Data, Tables 2-5). For the synthetic H3Y99-sulfopeptide, many nuclear protein tryptic peptides were observed. For example, histone H3 appeared with 82 peptide spectral matches (PSMs) and histone H4 has 8 PSMs, corresponding to 50% sequence coverage. **This significant carryover further calls into question the reported results as both a synthetic sulfo- and phosphopeptide were used by Yu et al**. We acquired Hep G2 whole cell lysate (WCL; Abcam ab166833) and nuclear extract lysate (NEL; ab14660). Both were digested with trypsin/Glu-C, followed by nanoLC-HCD MS/MS analysis. While we detected the AYLVGLFEDTNLC(carbamidomethyl)AIHAK peptide, there was no evidence of sulfation or phosphorylation. These samples, however, were not treated with phosphatase inhibitors. In addition, we performed Western blot analysis (anti-sulfotyrosine antibody; ab136481) of WCL and NEL with no evidence of histone sulfation (Extended Data, Figure 3).

Yu et al. proposed that SULT1B1 is responsible for histone H3 sulfation but their evidence again is based on LC-MS/MS data, which do not support this claim. The corresponding tryptic peptide MS/MS spectrum (Yu et al., Extended Data, Fig. 2g and **Figure 2a**) was acquired with ion trap collision induced dissociation (CID) at low resolution, which reduces confidence^3,9^. Sulfate-retaining fragments were annotated but, again, are not expected. Upon reexamination of this CID spectrum, we found some annotations questionable, including missing isotopic distributions, annotation of non-monoisotopic peaks, incorrect charge state assignment, and higher than typical error tolerance. We reannotated these data within 0.3 Da mass error, which removed all sulfate-retaining fragments (**Figure 2b**). Notably, this reannotation also showed a lack of fragments containing H3Y99, thus only confirming the N- and C-terminal ends of this larger tryptic peptide.

**Figure 2.**
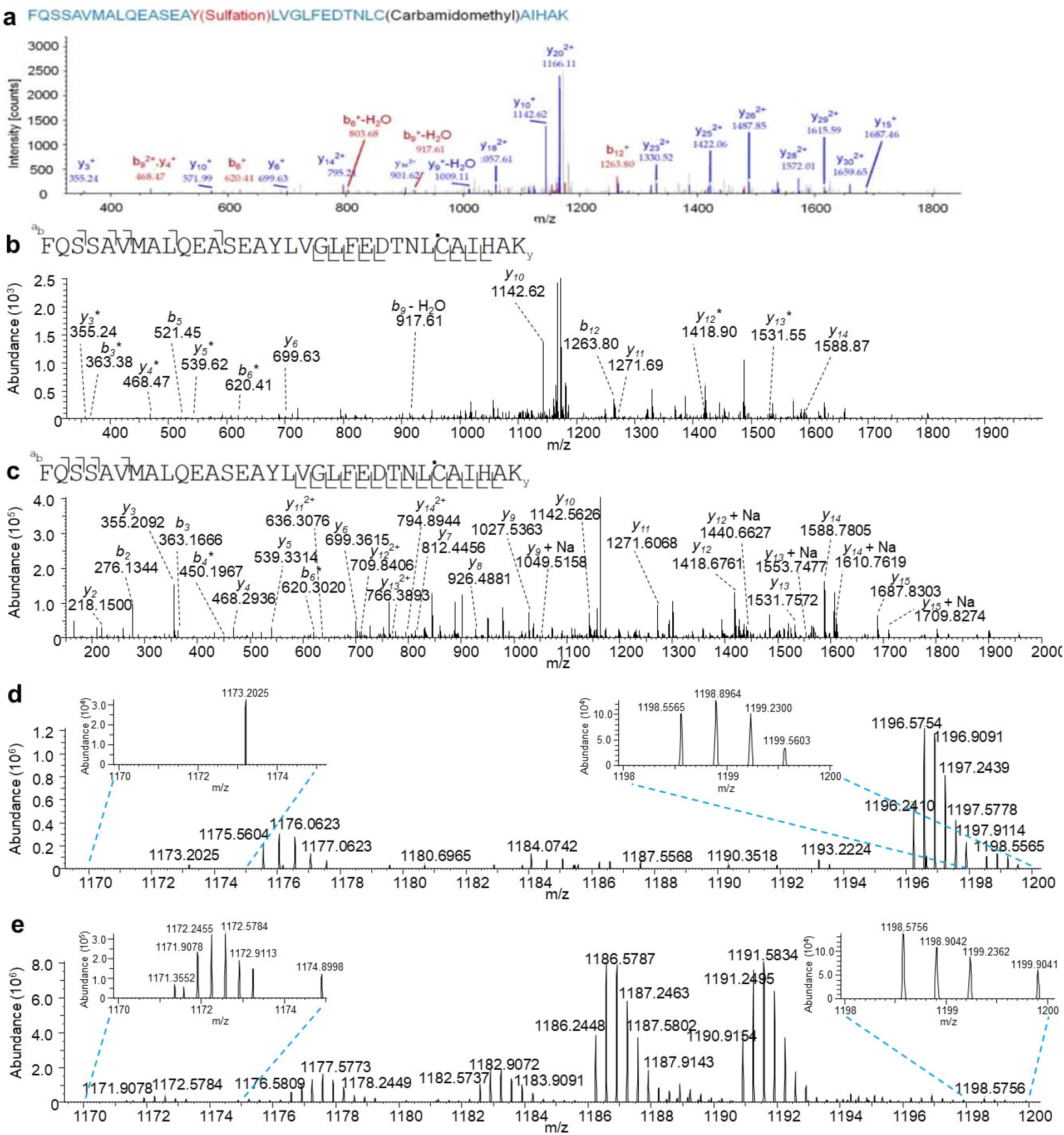
Lack of LC-MS/MS evidence supporting histone H3.2. Y99 sulfation upon incubation with SULT1B1 and PAPS. **a**, Low resolution CID MS/MS spectrum of the proposed, sulfated tryptic peptide FQSSAVMALQEASEAyLVGLFEDTNLC(carbamidomethyl)AIHAK, (y= sulfotyrosine), acquired and annotated by Yu et al. following incubation of histone H3.2. with SULT1B1 and PAPS. Several *y*-type ions containing the highly labile sulfate PTM were annotated. **b**, Reannotation of the CID spectrum in (a) within 0.3 Da mass accuracy. This reannotation removed all sulfate-retaining fragments and resulted in 45% sequence coverage with no information from the middle of the peptide, where the tyrosine residue is located. **c**, HCD MS/MS spectrum of histone H3.2. tryptic peptide FQSSAVMALQEASEAYLVGLFEDTNLC^▪^AIHAK (^▪^ = Carbamidomethylated cysteine) after subjecting histone H3.2. to PAPS and SULT1B1, showing an MS^1^ mass shift of ∼80 Da from the unmodified precursor ion. The corresponding modification site could not be determined due to lack of *y*-type fragments larger than *y15* and *b*-type fragments larger than *b*6 (all observed fragments lack a mass shift). Several sodium-adducted *y*-type fragments are noted, suggesting that such an alternative charge carrier may contribute towards the observed mass shift. **d**, MS^1^ spectrum of the proposed FQSSAVMALQEASEAyLVGLFEDTNLC(carbamidomethyl)AIHAK, (y= sulfotyrosine) peptide from Yu et al. showing the 3+ precursor ion at m/z 1198.5565 with no trace of the expected SO3 neutral loss at m/z 1171.9042. **e**, m/z 1198.5756 precursor ion (3+) observed in our MS^1^ data for the peptide FQSSAVMALQEASEAYLVGLFEDTNLC^▪^AIHAK (^▪^ = Carbamidomethylated cysteine). A corresponding ion (3+) lacking the sodium adduct is also seen at m/z 1191.2495 along with an isotopic distribution (3+) at m/z 1171.9078. However, the exact mass shift from the modified (presumably sodium-adducted) peptide precursor is - 80.0034 Da, 583 ppm off from an SO3 loss (79.9568 Da).

Furthermore, we subjected rh-histone H3.2 (catalog no. M2506S, New England BioLabs) to an rhSULT1B1 (R&D Systems) sulfation assay with PAPS (R&D Systems). This assay was optimized with the SULT1B1 substrate, 3,3′,5-triiodo-L-thyronine (T_3_, TCI America), showing a high degree of sulfation (**Extended Data, Figure 4**). By contrast, no evidence of rh-histone H3.2 sulfation was observed.

However; interestingly, a peptide with a mass similar to a potentially sulfated FQSSAVMALQEASEAyLVGLFEDTNLC(carbamidomethyl)AIHAK tryptic peptide was detected **(Figure 2c**). Similar to the results from Yu et al. information from the middle of the sequence was missing, hindering confident modification assignment and localization. Moreover, the signature 79.9568 SO_3_ neutral loss was not observed upon MS/MS (**Figures 2a-c**) or MS^1^ (**Figures 2d-e**), in either dataset.

Taken together, our new results along with reannotation of MS raw data from Yu et al. do not support the presence of histone H3 sulfation. Additionally, Yu et al. generated an anti-H3Y99sulf antibody, validated with the MODified Histone Peptide Array (catalog no.13001, Active Motif). However, this assay only validates N-terminal histone binding. Because Y99 is near the C-terminus, additional validation of this antibody seems warranted. Therefore, it is possible that the CUT&Tag results presented in Yu et al. may be due to phosphorylation rather than sulfation. Overall, our opinion is that the results and interpretations in Yu et al. warrant further validation and discussion, as they are both not clearly supported by the presented experimental data and contradictory to both literature precedent and our independent analyses.

## References

1. Lee, R.W. & Huttner, W.B. (Glu62, Ala30, Tyr8)n serves as high-affinity substrate for tyrosylprotein sulfotransferase: a Golgi enzyme. Proc. Natl. Acad. Sci. U. S. A. 82, 6143–6147 (1985).

2. Rossetto, D., Avvakumov, N. & Cote, J. Histone phosphorylation: a chromatin modification involved in diverse nuclear events. Epigenetics 7, 1098–1108 (2012).

3. Youssef, M.M. et al. Electron capture vs transfer dissociation for site determination of tryptic peptide tyrosine sulfation: direct detection of Fibrinogen sulfation sites and identification of novel isobaric interferences. J. Proteome Res. 23, 2386–2396 (2024).

4. Kweon, H.K. et al. Sulfoproteomics workflow with precursor ion accurate mass shift analysis reveals novel tyrosine sulfoproteins in the Golgi. J. Proteome Res. 23, 71–83 (2024).

5. Daly, L.A. et al. Custom workflow for the confident identification of sulfotyrosine-containing peptides and their discrimination from phosphopeptides. J. Proteome Res. 22, 3754–3772 (2023).

6. Nemeth-Cawley, J.F. et al. Analysis of sulfated peptides using positive electrospray ionization tandem mass spectrometry. J. Mass Spectrom. 36, 1301−1311 (2001).

7. Skulj, S. & Rozman, M. Study of the gas-phase fragmentation behaviour of sulfonated peptides. Int. J. Mass Spectrom. 391, 11−16 (2015).

8. Yu, et al. Determination of the sites of tyrosine O-sulfation in peptides and proteins. Nat. Methods 4, 583−588 (2007).

9. Ferries, S. et al. Evaluation of parameters for confident phosphorylation site localization using an orbitrap fusion tribrid mass Spectrometer. J. Proteome Res. 16, 3448–3459 (2017).

