## Supplemental Figures for "Established Sulfopeptide Tandem Mass Spectrometry Behavior and Sulfotransferase Assays Refute Tyrosine Sulfation as a Histone Mark"

**for**

Arising from W. Yu et al. *Nat. Chem. Biol*. **2023** (https://doi.org/10.1038/s41589-023-01267-9)

*Corresponding author.

**Extended Data Figure 1. Evidence of H3Y99 phosphopeptide presence in Yu et al. Hep G2 nuclear extract, mixed and synthetic sulfopeptide samples.**

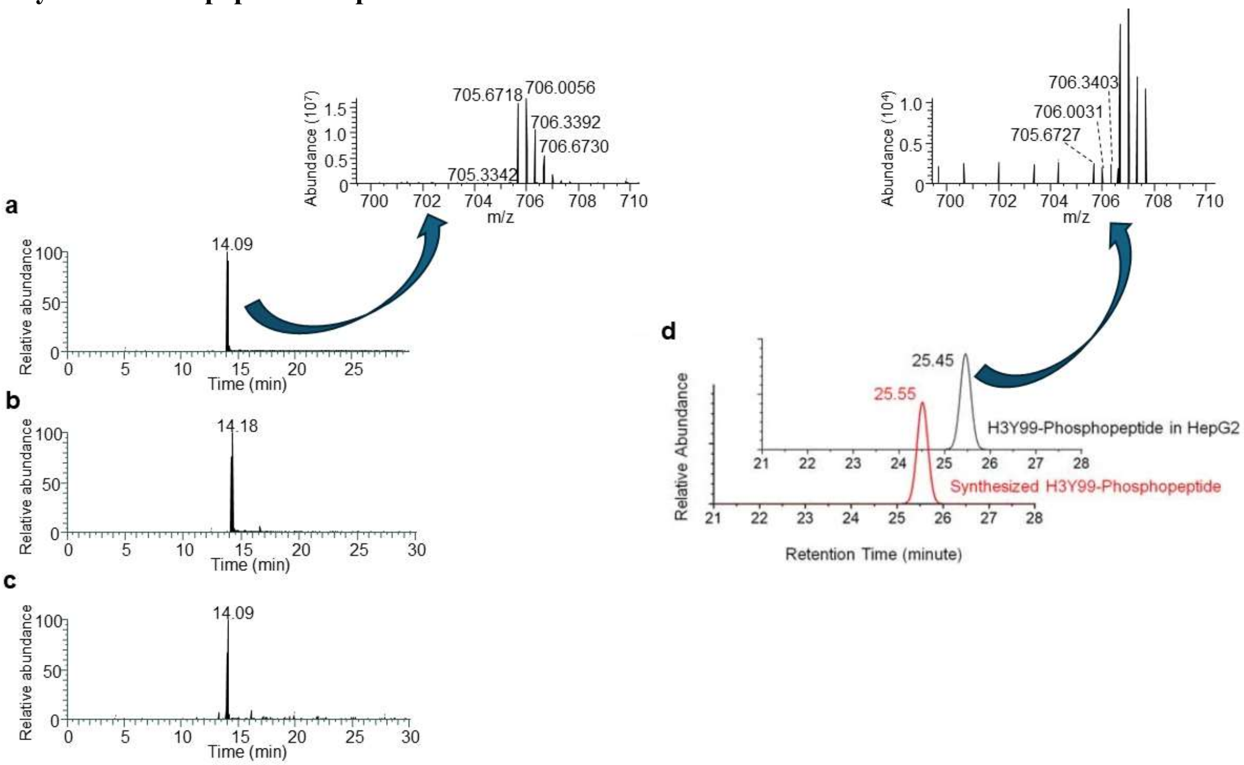


Extracted ion chromatograms of the tyrosine 99 phosphopeptide 3+ precursor ion (706.6709 m/z), at 6 ppm mass tolerance, in the HepG2 nuclear extract sample (**a**), mixed sample (**b**) and synthetic sulfopeptide sample (**c**) from Yu et al. The phosphopeptide was observed eluting at 14.09-14.18 min. in those short 30 min. gradient proteomic analysis runs. The corresponding $MS^1$ spectrum of the H3Y99 phosphopeptide precursor, eluting at 14.09 min in the HepG2 nuclear extract sample, showed 1.56e7 abundance (inset in **a**) compared to an abundance of 2.56e3 at 25.45 min (**d**) proposed by Yu et al. $MS^2$ spectrum supporting phosphopetide assignment was also absent around 25.45 min in Yu et al. Hep G2 sample. Moreover, no evidence of phosphopeptide was observed around this retention time in both Yu et al. mixed and synthetic sulfopeptide samples.

**Extended Data Figure 2. H3Y99 sulfation is misannotated in HCD MS/MS spectra from Yu et al. short proteomic runs.**

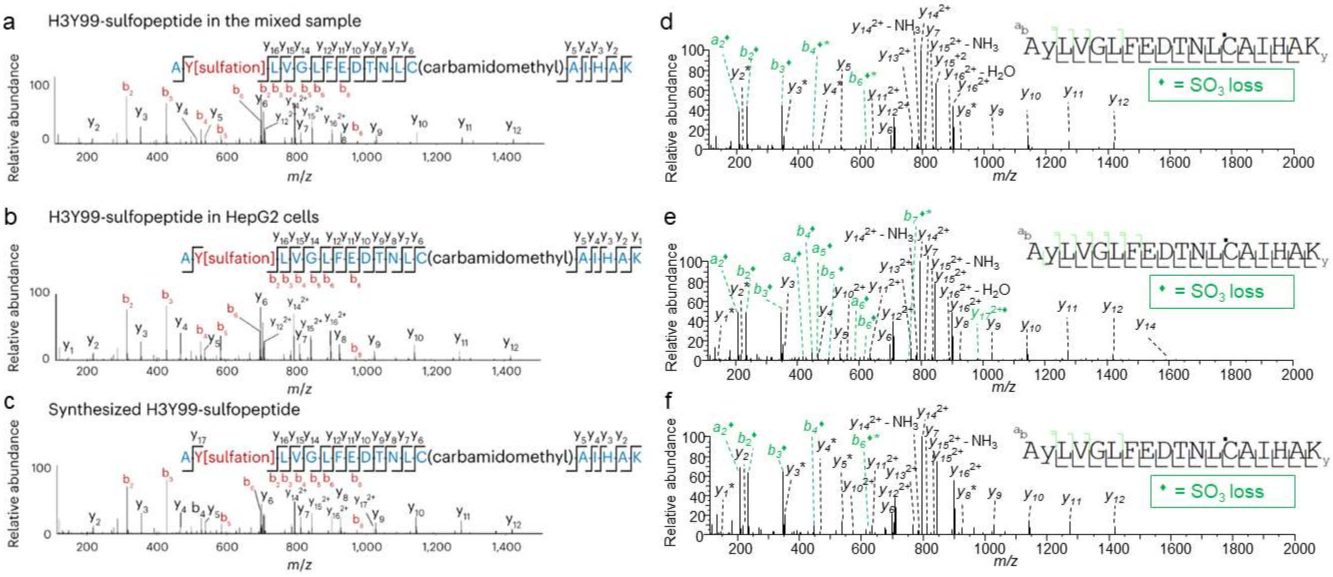


AyLVGLFEDTNLC(carbamidomethyl)AIHAK, (y= sulfotyrosine) HCD MS/MS spectra in the HepG2 nuclear extract sample (**a**), mixed sample (**b**) and synthetic sulfopeptide sample (**c**) as shown by Yu et al. The HCD MS/MS spectra annotations suggest *b*-type ions retaining the $SO_3$. Upon revisiting Yu et al.'s raw data, HCD MS/MS spectra showing a sulfopeptide fragmentation behavior with the complete $SO_3$ neutral loss were observed in the HepG2 nuclear extract sample (**d**), mixed sample (**e**) and synthetic sulfopeptide sample (**f**).

**Extended Data Figure 3. Undetected Histone H3 Tyrosine Sulfation in HepG2 Lysates.**

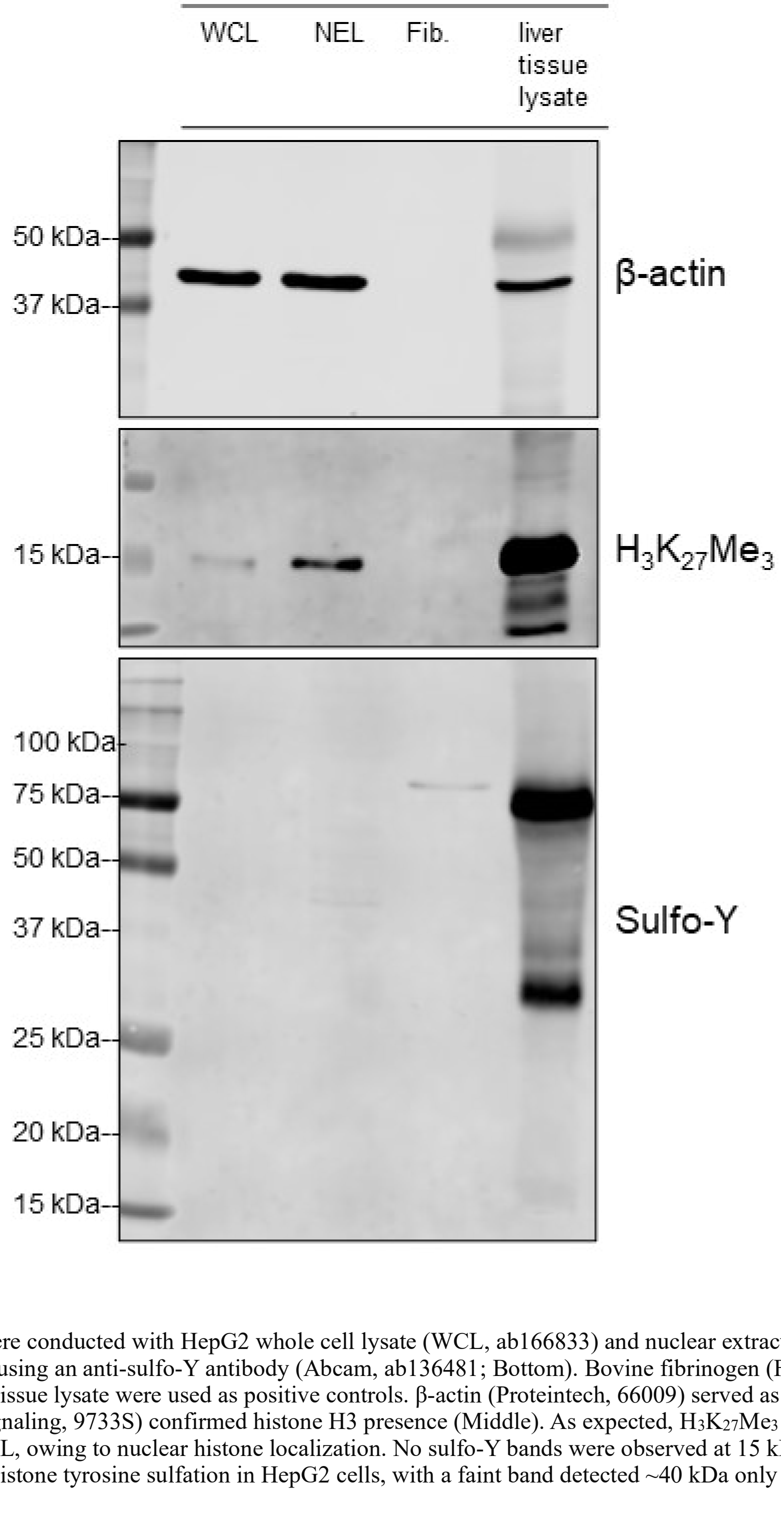


Western blot analyses were conducted with HepG2 whole cell lysate (WCL, ab166833) and nuclear extract lysate (NEL, ab14660) to detect tyrosine sulfation using an anti-sulfo-Y antibody (Abcam, ab136481; Bottom). Bovine fibrinogen (Fib., Sigma F8630, 0.5 µg/µl) and normal mouse liver tissue lysate were used as positive controls. β-actin (Proteintech, 66009) served as an internal control (Top). Anti-$H_3K_{27}Me_3$ (Cell Signaling, 9733S) confirmed histone H3 presence (Middle). As expected, $H_3K_{27}Me_3$ bands were more prominent in NEL compared to WCL, owing to nuclear histone localization. No sulfo-Y bands were observed at 15 kDa in either WCL or NEL, indicating insignificant histone tyrosine sulfation in HepG2 cells, with a faint band detected ~40 kDa only in NEL.

**Extended Data Figure 4. LC/ Fourier transform ion cyclotron resonance (FT-ICR) MS detection of sulfated T3, the known SULT1B1 substrate.**

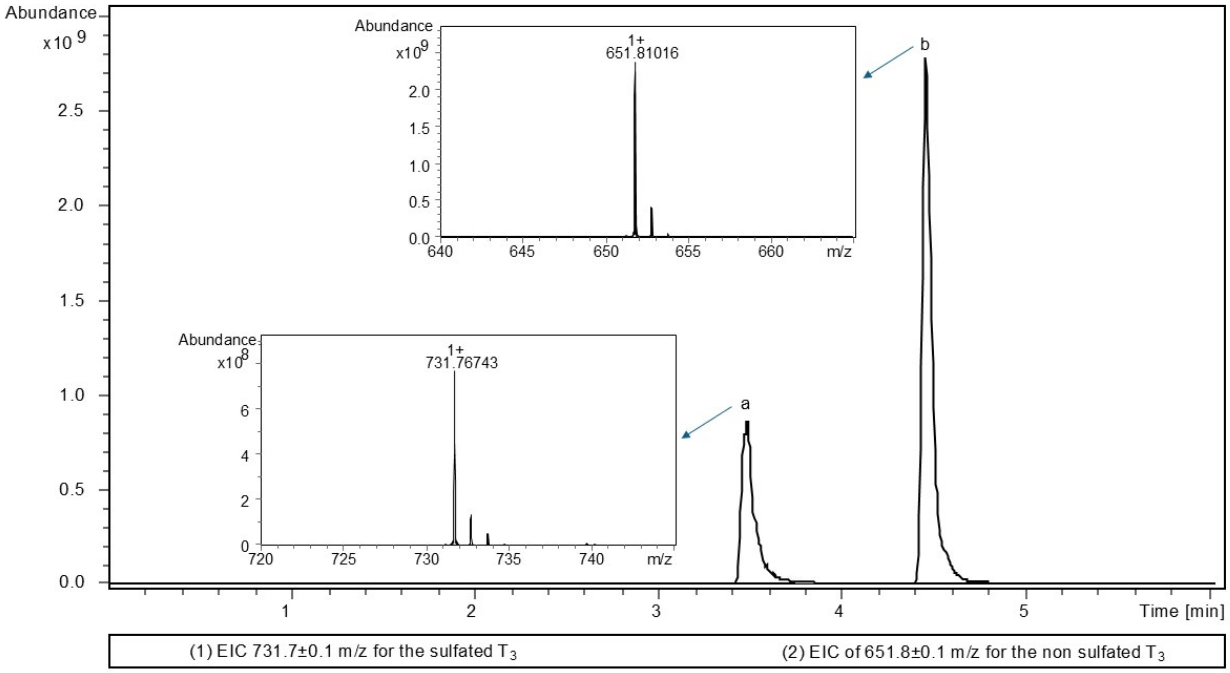


Activity of rhSULT1B1 was optimized using its known substrate, $T_3$, and LC/FT-ICR MS analysis achieved via positive polarity mode. Extracted ion chromatograms of sulfated $T_3$ (**a**) showed an abundance of 8e8, while non-sulfated $T_3$ (**b**) was at 2.5e9 abundance after incubation with rhSULT1B1 and PAPS at 37°C overnight.
